# Inhibiting BCKDK in triple negative breast cancer suppress protein translation, impair mitochondrial function, and potentiate doxorubicin cytotoxicity

**DOI:** 10.1101/2021.05.18.444634

**Authors:** Dipsikha Biswas, Logan Slade, Luke Duffley, Neil Mueller, Khoi Thien Dao, Angella Mercer, Yassine El Hiani, Petra Kienesberger, Thomas Pulinilkunnil

## Abstract

Triple-negative breast cancers (TNBCs) are characterized by poor survival, prognosis and gradual resistance to cytotoxic chemotherapeutics, like doxorubicin (DOX), which is limited by its cardiotoxic and chemoresistant effects that manifest over time. TNBC growth and survival are fuelled by reprogramming branched-chain amino acids (BCAAs) metabolism, which rewires oncogenic gene expression and cell signaling pathways. A regulatory kinase of the rate-limiting enzyme of the BCAA catabolic pathway, branched-chain ketoacid dehydrogenase kinase (BCKDK), have recently been implicated in driving tumor cell proliferation and conferring drug resistance by activating RAS/RAF/MEK/ERK signaling. However it remains unexplored if BCKDK remodels TNBC proliferation, survival and susceptibility to DOX-induced genotoxic stress. TNBC cell lines exhibited reduced BCKDK expression in response to DOX. Genetic and pharmacological inhibition of BCKDK in TNBC cell lines displayed reduced intracellular and secreted BCKAs. Moreover, BCKDK inhibition with concurrent DOX treatment exacerbated apoptosis, caspase activity and loss of TNBC proliferation. Transcriptome analysis of BCKDK silenced cells confirmed a marked upregulation of the apoptotic signaling pathway with increased protein ubiquitylation and compromised mitochondrial metabolism. BCKDK silencing in TNBC downregulated mitochondrial metabolism genes, reduced electron complex protein expression, oxygen consumption and ATP production. Silencing BCKDK in TNBC upregulated sestrin 2 and concurrently decreased protein synthesis and mTORC1 signaling. Inhibiting BCKDK in TNBC remodel BCAA flux, reduces protein translation triggering cell death, ATP insufficiency and susceptibility to genotoxic stress.

**Graphical abstract:** 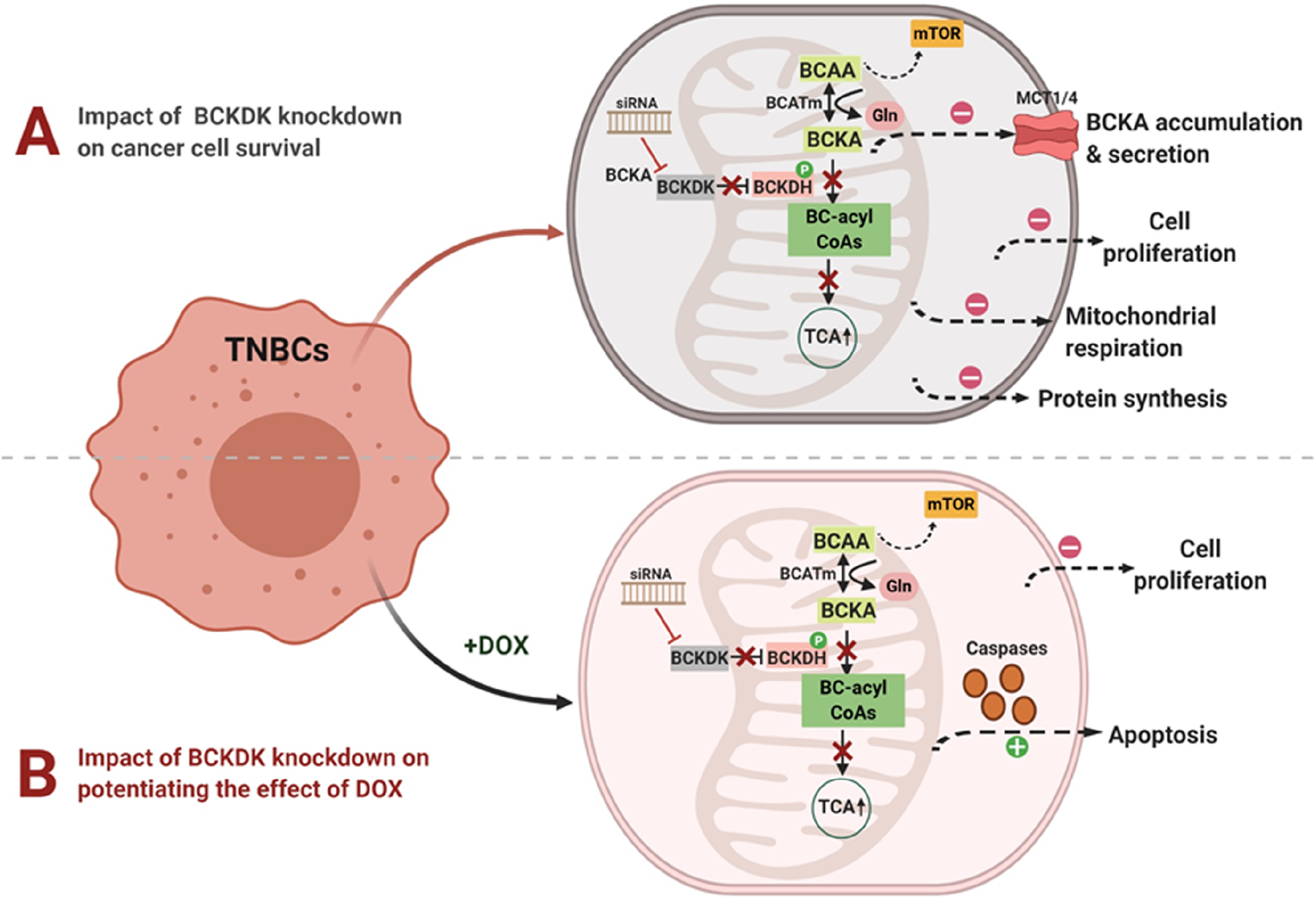

## Introduction

Triple-negative breast cancers (TNBCs) are an aggressive subtype of breast cancer, constituting 10-15% of all breast cancers and is defined by the lack of expression of estrogen receptor (ER), progesterone receptor (PR), and the human epidermal growth factor receptor 2 (HER2)^1^. Depending on the severity and stage of the disease^2^, TNBCs are generally treated in different combinations with genotoxic chemotherapies including anthracycline, taxane, docetaxel and cyclophosphamides. Doxorubicin (DOX) is an anthracycline class of cytotoxic chemotherapeutic that induces TNBC remission in about 30% of patients^3^; however, prolonged treatment with higher doses of DOX result in cardiotoxicity^4^, limiting its efficacy. Furthermore, breast cancer cells reprogram cellular metabolism and function to evade toxicity of chemotherapeutic agents, thereby exhibiting chemoresistance^5^. Metabolic maladaptation in TNBCs includes increased reliance on pentose phosphate pathway for NADPH, lysosomal and mitochondrial drug sequestration, altered nutrient metabolism, mitochondrial reprogramming, increased polyamine biosynthesis, glycolysis and glutaminolysis and amino acid uptake for protein biosynthesis^5^. Therefore, identifying mechanisms by which TNBCs survive DOX cytotoxicity and targeting the vulnerabilities in metabolic pathways that render TNBCs resistant to chemotherapy will enable developing concurrent therapies co-administered with lower doses of DOX.

Amino acid uptake and utilization facilitate the uninterrupted synthesis of TCA intermediates and proteins for increased nitrogen and proliferative demand of TNBCs^6^. Cancer cells tightly control the regulation of branched-chain amino acids (BCAAs), namely leucine, isoleucine and valine, which contribute to 20% of the TCA cycle intermediates. Besides direct incorporation into proteins, catabolism of BCAAs produces intermediates like glutamate vital for driving TNBC growth and survival^7^. BCAAs are reversibly transaminated to branched-chain ketoacids (BCKAs) by branched-chain aminotransferase (BCAT). BCKAs are further catabolized by branched-chain alpha-ketoacid dehydrogenase (BCKDH), ultimately generating TCA cycle intermediates. BCKDH is inhibited by phosphorylation at Ser 293 residue by the kinase, branched-chain alpha-ketoacid dehydrogenase kinase (BCKDK), while this phosphorylation is reversed by the Mg^2+^/Mn^2+^ Dependent 1K (PPM1K) protein phosphatase^7^. Recently in glioblastomas and pancreatic cancer, a role for BCKAs was uncovered in metabolic homeostasis^8^, immune suppression, and cell death evasion^9^. Indeed, degrading BCAAs and oxidizing BCKAs decreases cancer cell survival^10^. Although the role of BCKDH complex and BCAT isoforms (cytosolic BCAT1 or mitochondrial BCAT2) have been widely explored in different cancers^11^, the role of BCKDK remains mostly elusive. Recent studies have identified novel phosphorylation sites on BCKDK that stabilize BCKDK activity and influence tumor cell proliferation and survival^12,13^. Since reprogramming of BCAA metabolism produces intermediates that rewire oncogenic gene expression and cell signaling pathways in TNBC^11^, the importance of BCKDK in maintenance and utilization of BCAA pool, regulation of BCAA fate and BCAA flux into BCKA in TNBC growth and survival merits investigation.

In the current study, we investigated the role of BCKDK in TNBC survival and metabolism and examined whether DOX’s anticancer effect involves targeting the BCAA catabolic pathway. DOX treatment at 2μM suppressed BCKDK and significantly increased BCAA degradation enzyme mRNA and protein levels in two different TNBC cell lines. Transcriptome analysis of MDA-MB231 cells with BCKDK silencing revealed a marked increment in apoptotic signaling and deregulated mitochondrial function and biogenesis. Genetic and pharmacological inhibition of BCKDK sensitized TNBCs to DOX-induced cytotoxicity. BCKDK inhibition suppressed mitochondrial function, reduced nascent protein synthesis and increased sestrin 2 (SESN2) expression, a negative regulator of protein synthesis and cell proliferation. Collectively, our data highlight the role of BCKDK as a novel DOX sensitive target in TNBCs. Targeting BCKDK can be an attractive therapeutic target in regulating TNBC survival, proliferation, and potentiating chemosensitivity to DOX.

## Results

### Altered BCAA catabolic enzyme expression in TNBCs is sensitive to DOX treatment

BCAA catabolic enzyme expression was examined in two TNBC cell lines, BT549 and MDA-MB231. In agreement with prior studies^14^, we found a striking upregulation of *BCAT1* mRNA levels, specifically in MDA-MB231 cells, while *BCAT2* mRNA (Fig 1A) and protein (Fig S1B) levels were significantly downregulated in both BT549 and MDA-MB231 cells when compared with normal breast epithelial cell line, MCF10A. Treatment with 2μM DOX reduced *BCAT1* and further reduced *BCAT2* mRNA expression in both TNBCs (Fig S1C-D). BCKDHA and BCKDHB protein levels were significantly downregulated in TNBC cell lines (Fig S1B), although their mRNA levels were selectively reduced in the MDA-MB231 cells (Fig 1A). DOX treatment increased *BCKDHA* mRNA (Fig 1B) and protein (Fig 1D) levels in MDA-MB231 cells. In the TNBC cells, mRNA (Fig S1A) and protein (Fig S1B) expression of *KLF15*, the transcriptional activator of the BCAA pathway, was significantly suppressed. DOX treatment significantly increased *KLF15* mRNA (Fig 1B-C) expression in both TNBCs and KLF15 protein content specifically (Fig 1D) in MDA-MB231 cells. The BCKDH phosphatase, *PPM1K*, mRNA levels were also decreased in TNBCs (Fig 1A) and were upregulated in response to DOX (Fig 1B-C). mRNA expression of distal enzymes, *ACADSB*, *HADHA*, *HIBCH* were unchanged (Fig S1A) and DOX increased the mRNA expression of *HADHA* in both TNBCs (Fig S1C-D). These data suggest that in TNBCs BCAA catabolic enzyme expression is suppressed, and treatment with DOX counteracts this suppression, likely affecting BCAA catabolism. Since the role of BCAT and BCKDH were examined in prior studies, we focussed on studying the role of BCKDK, the kinase which regulates the flux of BCKAs towards oxidation by inactive phosphorylation of BCKDH. *BCKDK* transcripts were markedly upregulated in BT549 and marginally increased in MDA-MB231 cells (Fig 1A). Although BCKDK protein expression was augmented in BT549, it remained comparable (p=0.08) to MCF10A cells in MDA-MB231 cells (Fig S1B). The phosphorylated to total BCKDE1α S293 content was increased in BT549 but not in MDA-MB231 and MCF10A cells (Fig S1B). DOX treatment reduced *BCKDK* mRNA levels in both TNBCs (Fig 1B-C). 2μM DOX treatment reduced BCKDK protein levels and the corresponding phosphorylated BCKDE1α S293 levels in MDA-MB231 (Fig 1D) and BT549 cells (Fig S1E). Since BCKDK expression was altered in TNBCs and regulated by DOX, we next examined the significance of BCKDK in TNBC metabolism and proliferation.

**Figure 1.**
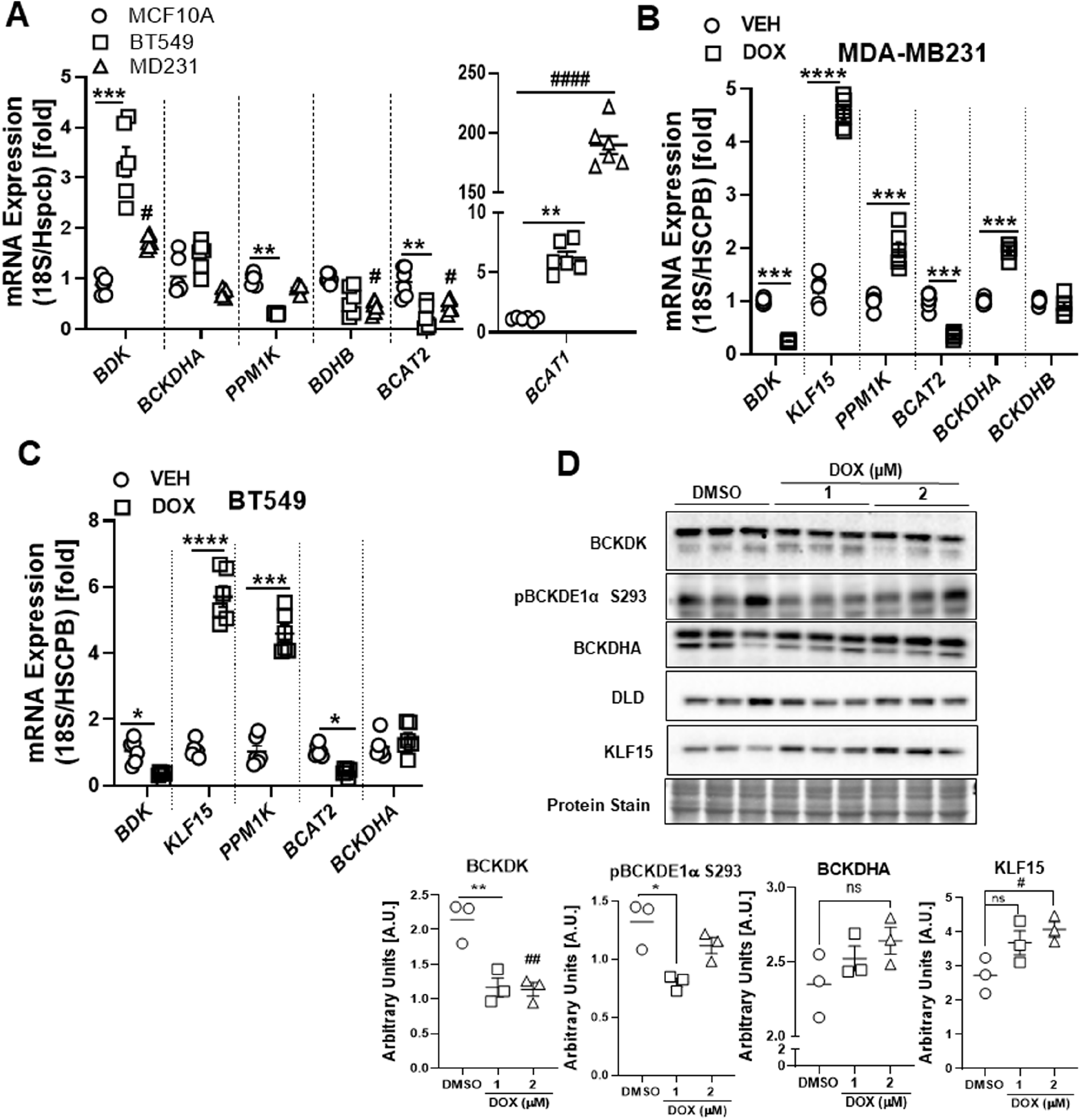
DOX suppresses BCKDK expression and augments BCAA oxidation enzyme expression in TNBCs. A) Quantification of *BCKDK*, *BCKDHA*, *PPM1K*, *BCKDHB*, *BCAT2*, and *BCAT1* BCAA catabolic enzyme mRNA expression corrected to 18S/HSPCB reference genes in MCF10A, BT549 and MDA-MB231 cells. B-C) BCAA catabolic enzyme expression in TNBCs treated with 2μM DOX for 18h. Quantification of *BCKDK*, *BCKDHA*, *PPM1K*, *BCKDHB*, *BCAT2*, and *KLF15* mRNA expression corrected to 18S/HSPCB reference genes in MDA-MB231 (B) and BT549 (C) cells. D) Immunoblot and densitometric analysis of BCKDK, total and phosphorylated BCKDHA E1α Ser 293, DLD and KLF15 in MDA-MB231 cells treated with 1μM and 2μM DOX for 18h. Quantifications are from three independent experiments. Data presented as mean ± S.D. Statistical analysis was performed using a two-way ANOVA followed by a Tukey’s multiple comparison test; *p <0.05, **p < 0.01, **** p <0.0001 as indicated. *MCF10A v/s BT549 or DMSO v/s 1μM DOX, #MCF10A v/s MDA-MB231 or DMSO v/s 2μM DOX.

### Downregulating BCKDK inhibits proliferation, promotes apoptosis and potentiates DOX mediated cell death

To decipher whether BCKDK silencing alters metabolism and survival of TNBCs, MDA-MB231 cells were treated with either control siRNA (siCON) or siRNAs targeting BCKDK exon 2 (siBDK#1) or exon 7 (siBDK#2). BCKDK silencing was confirmed by decreased mRNA (Fig S2A) and protein (Fig S2B) levels. Silencing BCKDK using siBDK#2 resulted in reduced cell count at both 24h and 48h following transfection, while siBDK#1 showed reduced cell count at 48h (Fig S2C), suggesting that targeting BCKDK influences TNBC survival and proliferation. BCKDK deletion also resulted in significant upregulation of cleaved caspase 3 content (Fig S2B), indicating increased cell death. In an antibody array of apoptosis-associated proteins, levels of the anti-apoptotic protein, survivin were decreased and pro-apoptotic protein, BIM and cytochrome C levels were increased in BCKDK silenced MDA-MB231 cells while potentiating DOX mediated cytochrome C, Fas, BIM and caspase 8 expression (Fig 2A). Further, BCKDK knockdown exacerbated DOX mediated increases in cleaved caspase 3, caspase 7 and PARP levels in MDA-MB231 cells (Fig 2B). In BCKDK depleted cells, exposure to DOX increased phosphorylation of the DNA damage marker, ataxia-telangiectasia mutated (ATM), suggesting increased genomic instability and apoptosis (Fig 2B). Similar increases was observed in cleaved caspase 3/7 and BAX Bax expression in DOX-treated BT549 cells with silencing BCKDK (Fig S2D). To rule out off-target effects of siRNA mediated gene silencing, we also performed the experiments with adenoviral-mediated knockdown of BCKDK in both MDA-MB231 and BT549 cells and observed similar increases in cleaved caspase levels in response to DOX (Fig S2E). Changes in protein levels were also reflected in increased caspase activity in BCKDK depleted MDA-MB231 (Fig 2C) and BT549 (Fig S2F) cells treated with DOX. Cytotoxicity of DOX was evident by increased lactate dehydrogenase (LDH) release into the media, which was exacerbated further following BCKDK silencing (Fig 2D).

**Figure 2.**
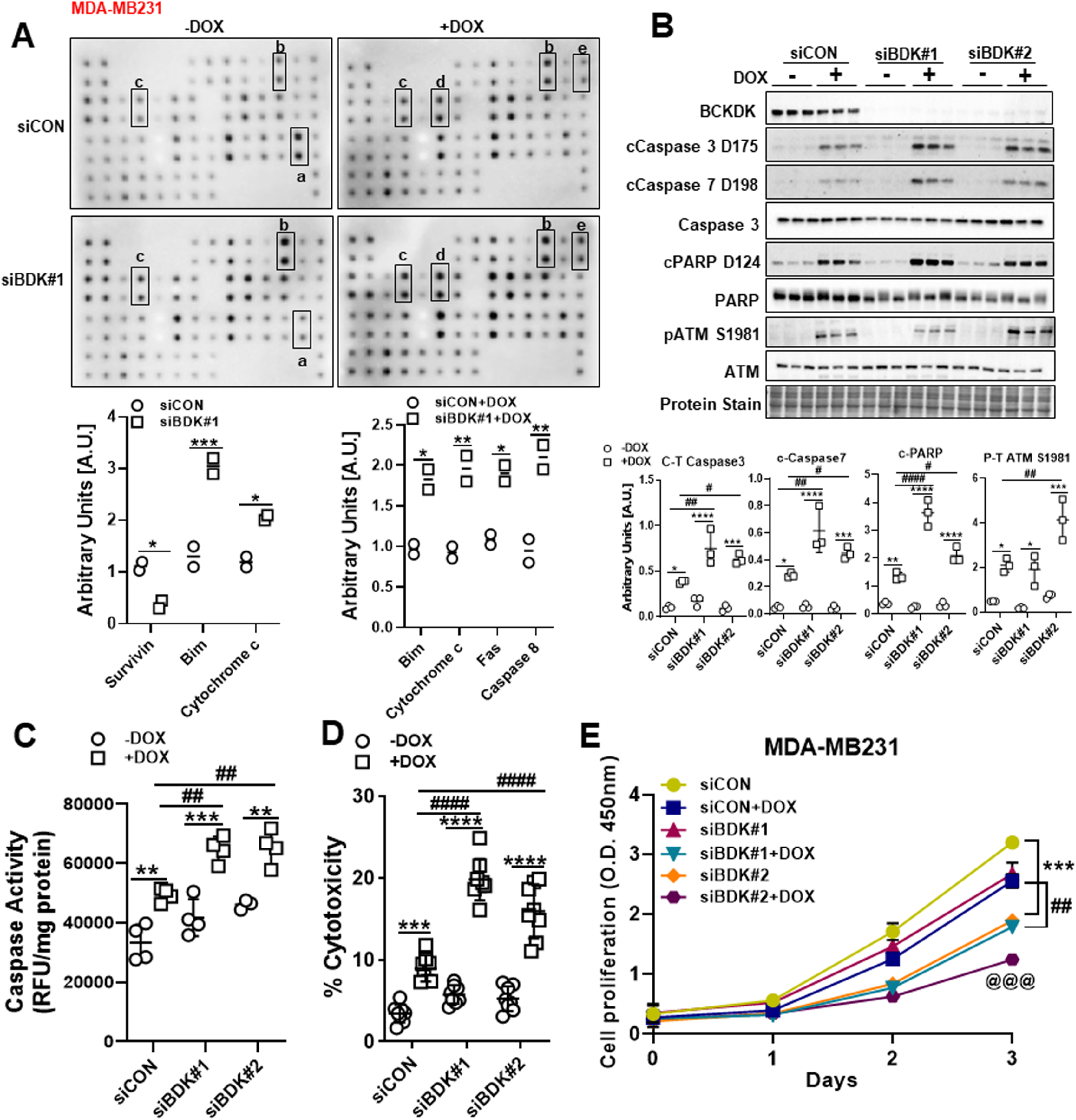
Silencing BCKDK exacerbates DOX mediated cell death in TNBCs. A) Antibody-immobilized PVDF membranes containing 43 different apoptosis associated proteins incubated with extracts of CON (siCON) and BCKDK (siBDK#1) silenced MDA-MB231 cells, treated with or without 2μM DOX for 18h. Differentially expressed proteins (fold change more than 1.5 or less than 0.6) are quantified and indicated on the blots. a=Survivin, b=Bim, c=Cytochrome, d=Fas, e= Caspase8. B) Immunoblot and densitometric analysis of BCKDK, total and cleaved Caspase 3, cleaved Caspase 7, total and cleaved PARP, total and phosphorylated ATM Ser 1981 in MDA-MB231 cells transfected with siCON, siBDK#1 or siBDK#2 for 72h followed by 2μM DOX or DMSO for 18h. (C-E) MDA-MB231 cells were transfected with siCON, siBDK#1 or siBDK#2 for 72h followed by 2μM DOX or DMSO treatment for 18h and analyzed for C) Caspase 3 activity, D) LDH release in the media, and E) cell proliferation measured by CCK8 assay. Data presented as mean ± S.D. Statistical analysis was performed using a two-way ANOVA followed by a Tukey’s multiple comparison test; *p <0.05, **p < 0.01, **** p <0.0001 as indicated.

To examine if the increased cytotoxic response of MDA-MB231 cells to DOX following BCKDK knockdown was associated with decreased proliferation, CCK-8 assay was performed. BCKDK knockdown alone reduced cell proliferation by 2d (siBDK#2) or 3d (siBDK#1) which was potentiated further by DOX treatment (Fig 2E). Additionally, MDA-MB231 cells transduced with shBCKDK were treated with DOX for 18h followed by a chase in drug-free media for 48h. Cell viability was determined by using the Presto Blue reagent wherein reduction in resazurin is the surrogate readout for metabolic activity. Knockdown of BCKDK resulted in a significant decrease in metabolic activity that further declined by exposure to DOX (Fig S2G).

We next recapitulated the cytotoxic effects of BCKDK silencing by treating cells with the BCKDK inhibitor, BT2, in the presence and absence of DOX. BT2 treatment alone resulted in increased levels of cleaved PARP, caspase 3 and caspase 7 and phosphorylated ATM in MDA-MB231 cells (Fig 3A). BT2 also potentiated DOX’s effects on cleaved caspase3, cleaved PARP and pATM levels in BT549 cells (Fig 3B), with the maximal effects observed at 500 μM. BT2 markedly exacerbated DOX mediated increase in caspase activity in MDA-MB231 (Fig 3C) and BT549 (Fig 3D) cells. Moreover, in MDA-MB231 cells BT2 alone increased LDH release, and this effect was further exacerbated in the presence of DOX (Fig 3E).

**Figure 3.**
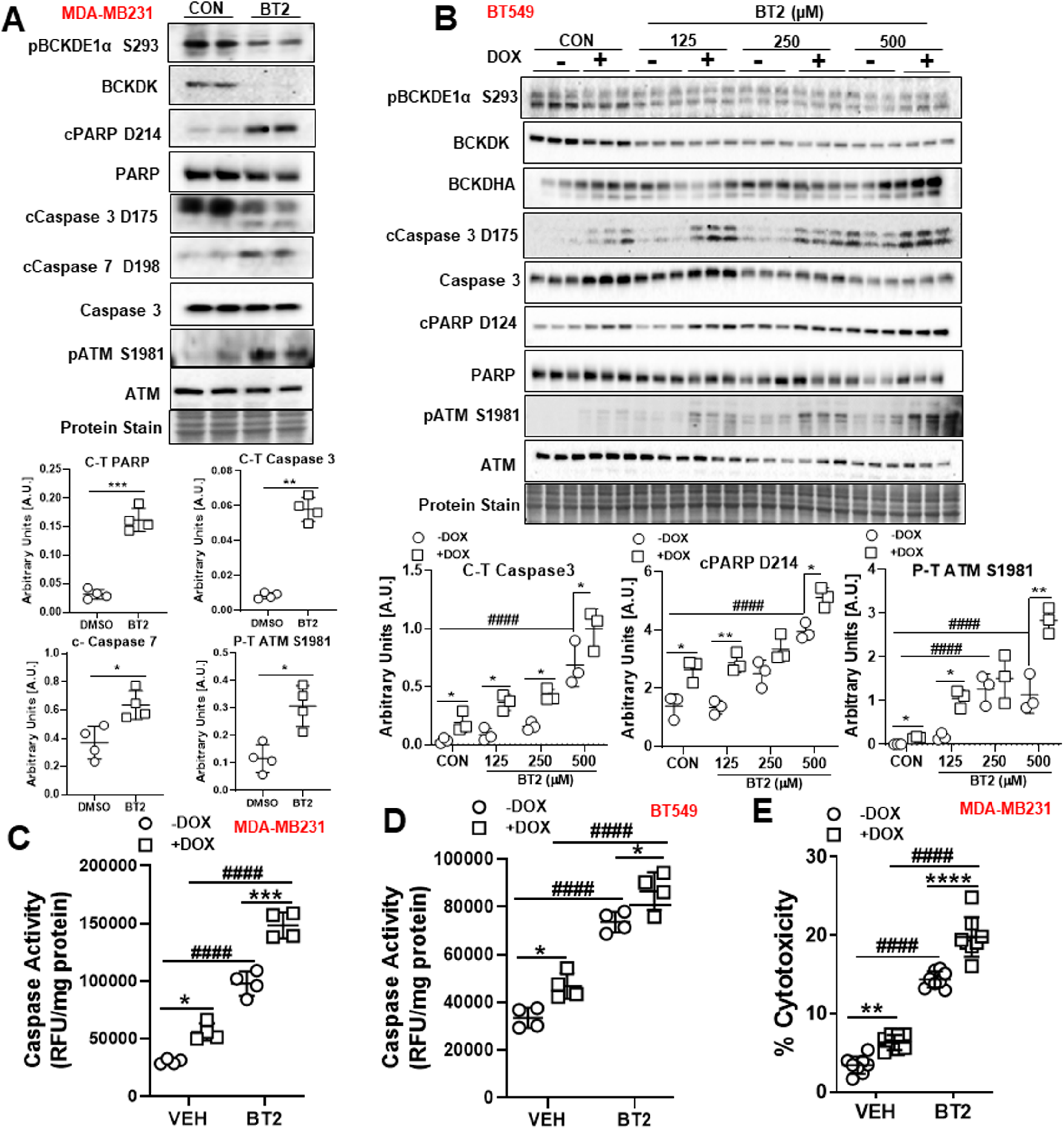
BT2 mediated BCKDK inhibition induces cell death and potentiates DOX mediated apoptosis in TNBCs. A) Immunoblot and densitometric analysis of phosphorylated BCKDE1α Ser 293, BCKDK, total and cleaved Caspase 3, cleaved Caspase 7, total and cleaved PARP, total and phosphorylated ATM Ser 1981 in MDA-MB231 cells treated with 500μM BT2 for 20h. Densitometric analysis is from three independent experiments. B) Immunoblot and densitometric analysis of total and phosphorylated BCKDE1α Ser 293, BCKDK, total and cleaved Caspase 3, total and cleaved PARP, total and phosphorylated ATM Ser 1981 in MDA-MB231 cells pre-treated with 500μM BT2 for 20h followed by 2μM DOX or DMSO treatment for 18h. Caspase 3 activity measured in MDA-MB231 (C) and BT549 (D) cells and LDH release into the media was measured in MDA-MB231 cells (E) pre-treated with 500μM BT2 for 20h followed by 2μM DOX or DMSO treatment for 18h. Quantifications are from three independent experiments. Data presented as mean ± S.D. Statistical analysis was performed using a two-way ANOVA followed by a Tukey’s multiple comparison test; *p <0.05, **p < 0.01, **** p <0.0001 as indicated.

### Silencing BCKDK alters TNBC transcriptome

We next aimed to identify metabolic and signaling pathway networks regulated by BCKDK to influence TNBC cell death and proliferation. MDA-MB231 cells depleted of BCKDK were subjected to RNA-Seq transcriptomic analysis. Principle component analysis and distance clustering showed that the transcriptome of cells treated with siBDK#1 and siBDK#2 displayed little similarity (Fig 4A-B), thus we classified genes as differentially expressed compared with control if the adjusted P-value was below 0.01 for each siRNA individually and the fold change was occurring in the same direction. This method identified 1024 genes, which were differentially expressed by both BCKDK siRNA’s and over-represented Gene Ontology (GO) were determined using DAVID (Fig 4C). Among the top 20 GO terms identified, activation of response to unfolded protein response (UPR) and positive regulation of ubiquitin-dependent protein catabolism appeared multiple times (Fig 4D). Both these processes play a crucial role in regulating apoptosis by direct or indirect targeting of important regulators of apoptosis and caspases^15,16^. Indeed, the GO term related to activation of caspases, the cysteine-type endopeptidase, was enriched in the differentially expressed genes from BCKDK knockdown cells (Fig 2D), consistent with our data (Fig 2-3). Genes annotated to this GO term include the initiator or activator caspases, including CASP2, CASP8, CASP9, and CASP10, also called initiator (or apical, or activator) caspases, suggesting that BCKDK silencing is plausibly sensitizing TNBC to cell death. Indeed, differentially expressed genes involved in death receptor signaling was upregulated after BCKDK knockdown, including TNF-related apoptosis-inducing ligand (TRAIL/TNFSF10) and TNFRSF1A associated via death domain (TRADD) (Fig 4E). The expression of pro-apoptotic factor, Bax and Smad3, involved in activating TGF-β-induced apoptosis, increased in BCKDK depleted cells (Fig 4E), indicating augmented pro-apoptotic gene expression.

**Figure 4.**
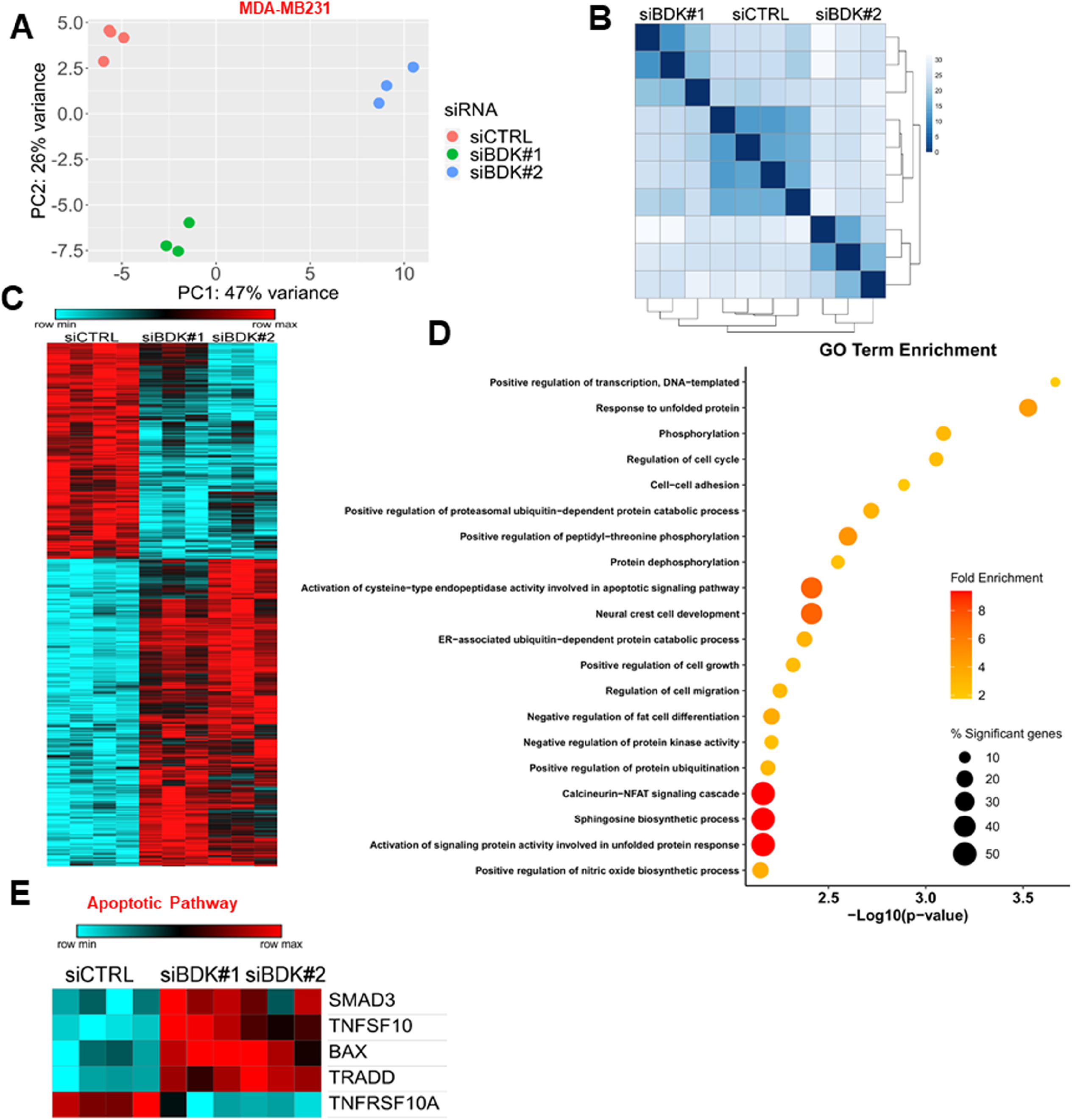
Differential enrichment of genes related to metabolism and cell death pathways in MDA-MB-231 cells with BCKDK knockdown. A) Principle component analysis plot for all gene expression from MDA-MB231 cells treated with both siRNAs targeting BCKDK. B) Distance matrix clustering for all gene expression from MDA-MB231 cells. C) Heatmap for differentially expressed genes from MDA-MB231 cells. D) Top 20 most significantly enriched gene ontology (GO) terms from BCKDK knockdown induced differentially expressed genes, color represents the fold enrichment statistic, and size represents the percentage of the differentially expressed genes in the gene set compared with the total gene set size. E) Heatmap for genes involved in apoptosis which were significantly differentially regulated by BCKDK knockdown in MDA-MB231 cells.

### BCKDK inhibition decreases intracellular and secreted BCKAs

We next ascertained the consequences of BCKDK silencing in influencing BCKA flux by measuring mRNA expression of BCAA catabolic enzymes that facilitate BCKA oxidation. RNA-Seq analysis revealed that BCKDK knockdown resulted in increased expression of genes involved in BCKA oxidation, specifically *BCKDHA*, methylmalonyl-CoA epimerase (*MCEE*), propionyl-CoA carboxylase (*PCCB*), *HADHB* and *ACADM* (Fig S3A). BCKDK depleted cells also displayed reduced expression of *BCAT1* and *HIBCH* (Fig S3A), genes augmented in several cancers^17^. qPCR analysis revealed significant increases in the BCKDHA and KLF15 mRNA expression in MDA-MB231 (Fig S3B), although it was only observed with siBDK#1. No changes were observed in the mRNA expression of BCAA catabolic enzymes in the BT549 cells (Fig S3C). Contrary to BCKDK silencing, BT2 treatment resulted in significant increases in *KLF15*, *BCKDHA*, *BCKDHB*, *HIBCH* and *HIBADH* mRNA levels in BT549 (Fig S3D). We next ascertained whether increased expression of BCAA degradation genes resulted in reduced intracellular accumulation and secretion of BCKAs. Adenoviral knockdown of BCKDK in MDA-MB231 cells resulted in decreased intracellular BCKAs accumulation as revealed by UPLC-MS/MS analysis (Fig S3E). A significant reduction in total BCKAs was observed in the cells and in the media in MDA-MB231 cells treated with BT2 (Fig S3F-G). Previous studies have demonstrated the role of tumor secreted BCKAs in proliferation and immune suppression of cancer cells^8,9^. Our data suggest that the mechanism by which BCKDK depletion arrests TNBC proliferation likely involves increased BCKA oxidation with a concomitant reduction in accumulation and secretion of BCKAs.

### BCKDK inhibition results in decreased mitochondrial complex protein expression and impaired mitochondrial function

We next ascertained if increased cell death following BCKDK silencing is an outcome of mitochondrial dysfunction. Indeed, permeabilized and damaged mitochondria generate reactive oxygen species in a complex I and II-dependent manner^18^ and when recognized by activated caspase is targeted for destruction. Transcriptomic analysis of BCKDK depleted MDA-MB231 cells revealed a marked downregulation of genes involved in mitochondrial function and biogenesis (Fig 5A). For instance, BCKDK silencing reduced nucleoside diphosphate kinase (NME4) and clusterin (Clu) gene expression, which are involved in promoting redistribution of cardiolipin to prevent mitochondrial permeabilization and suppressing Bax dependent release of cytochrome C, respectively (Fig 5A). Genes involved in mitochondrial translation, mitochondrial ribosome construction and abundance of 16S mt-rRNA, such as G elongation factor mitochondrial 1 (GFMI), mitochondrial ribosomal protein L19 (MRPL19) and RNA pseudouridine synthase D4 (RPUSD4), were also downregulated upon BCKDK silencing (Fig 5A). Furthermore, genes involved in complex I assembly (NADH:ubiquinone oxidoreductase assembly factor 1, NDUFAF1), the formation of complex III (tetratricopeptide repeat domain 19, TTC19), suppression of ROS (superoxide dismutase 2, SOD2) and synthesis of nucleotide triphosphates and mitochondrial respiration (NME4, serine active site containing 1, SERAC) were downregulated in BCKDK depleted cells (Fig 6A). Additionally, mitochondrial genes involved in pyruvate transport (mitochondrial pyruvate carrier 1, MPC1), glycine metabolism (glycine cleavage system protein H, GCSH and glycine c-acetyltransferase, GCAT) and ether lipid synthesis and intracellular cholesterol trafficking (peroxisomal alkylglycerone phosphate synthase, AGPS and mitochondrial SERAC1, respectively) are also downregulated upon BCKDK silencing (Fig 5A). Consistent with the transcriptomic data, silencing BCKDK in MDA-MB231 cells resulted in decreased complex I, complex II and complex III protein levels (Fig 5B). Extracellular flux analysis studies revealed reduced basal oxygen consumption rates (OCR) (Fig 5C), ATP production (Fig 5D) and non-mitochondrial respiration (Fig S4A) in the BCKDK silenced cells. Inhibiting BCKDK by BT2 also resulted in reduced basal and maximal OCR (Fig 5E) and ATP production (Fig 5F), indicating impaired mitochondrial oxidative metabolism. Lower OCR also resulted in reduced spare respiratory capacity (Fig 5G) and proton leak (Fig 5H) as well as low non-mitochondrial respiration (Fig S4B), suggestive of impaired adaptation to metabolic changes. Mitochondrial DOX accumulation can also disrupt mitochondrial architecture and oxidative capacity^19^. We determined whether BCKDK silencing in MDA-MB231 cells can potentiate DOX’s effects, exacerbating mitochondrial dysfunction. DOX treatment reduced basal OCR (Fig 5C), ATP production (Fig 5D) and non-mitochondrial respiration (Fig S4A), but it was not significantly potentiated upon BCKDK silencing. Unlike TNBCs, normal breast epithelial cells, MCF10A, when treated with BT2, did not show suppressed mitochondrial respiration or ATP production (Fig S4 C-G).

**Figure 5.**
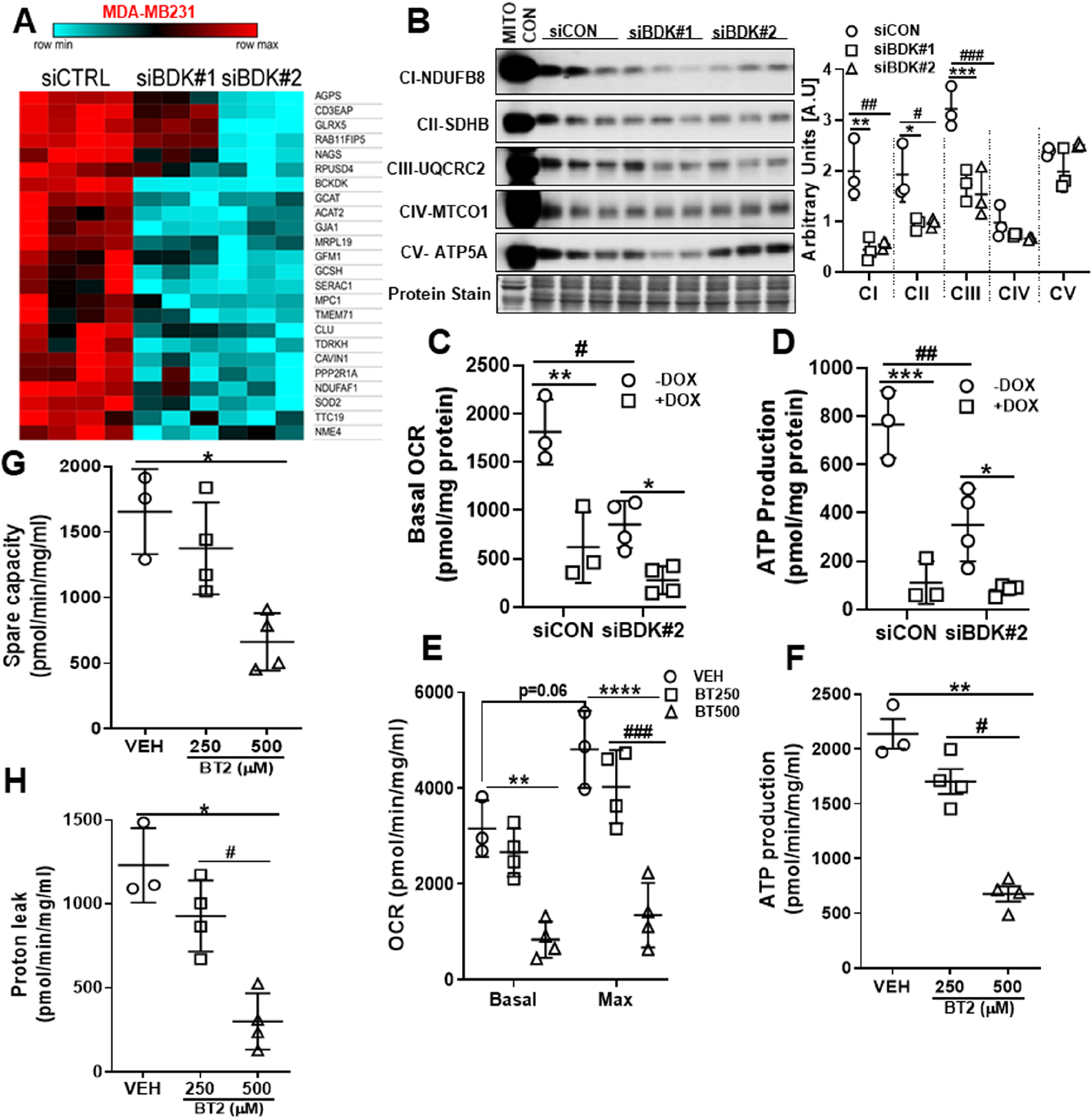
BCKDK inactivation suppresses mitochondrial function and complex protein expression. A) Heatmap for genes involved in mitochondrial biogenesis, structure, and function was significantly differentially regulated by BCKDK knockdown. B) Immunoblot and densitometric quantification of complex I-V proteins in MDA-MB231 whole cell lysates transfected with siCON, siBDK#1 or siBDK#2 for 72h. Data presented as mean ± S.D. Statistical analysis was performed using a two-way ANOVA followed by a Tukey’s multiple comparison test;*p <0.05, **p < 0.01, **** p <0.0001 as indicated. (C-H) Mitochondrial function measured in BCKDK depleted MDA-MB231 cells using extracellular flux analyzer in the presence of 25mM glucose. C) Basal OCR, D) ATP production measured in MDA-MB231 cells transfected with siCON or siBDK#2. E) Basal and maximal OCR, F) ATP production, G) spare capacity, and H) proton leak measured in MDA-MB231 cells treated with 250μM or 500μM BT2 for 20h. Data presented as mean ± S.D. Statistical analysis was performed using a was performed using Student’s t-test; *p <0.05, **p < 0.01, **** p <0.0001 as indicated.

**Figure 6.**
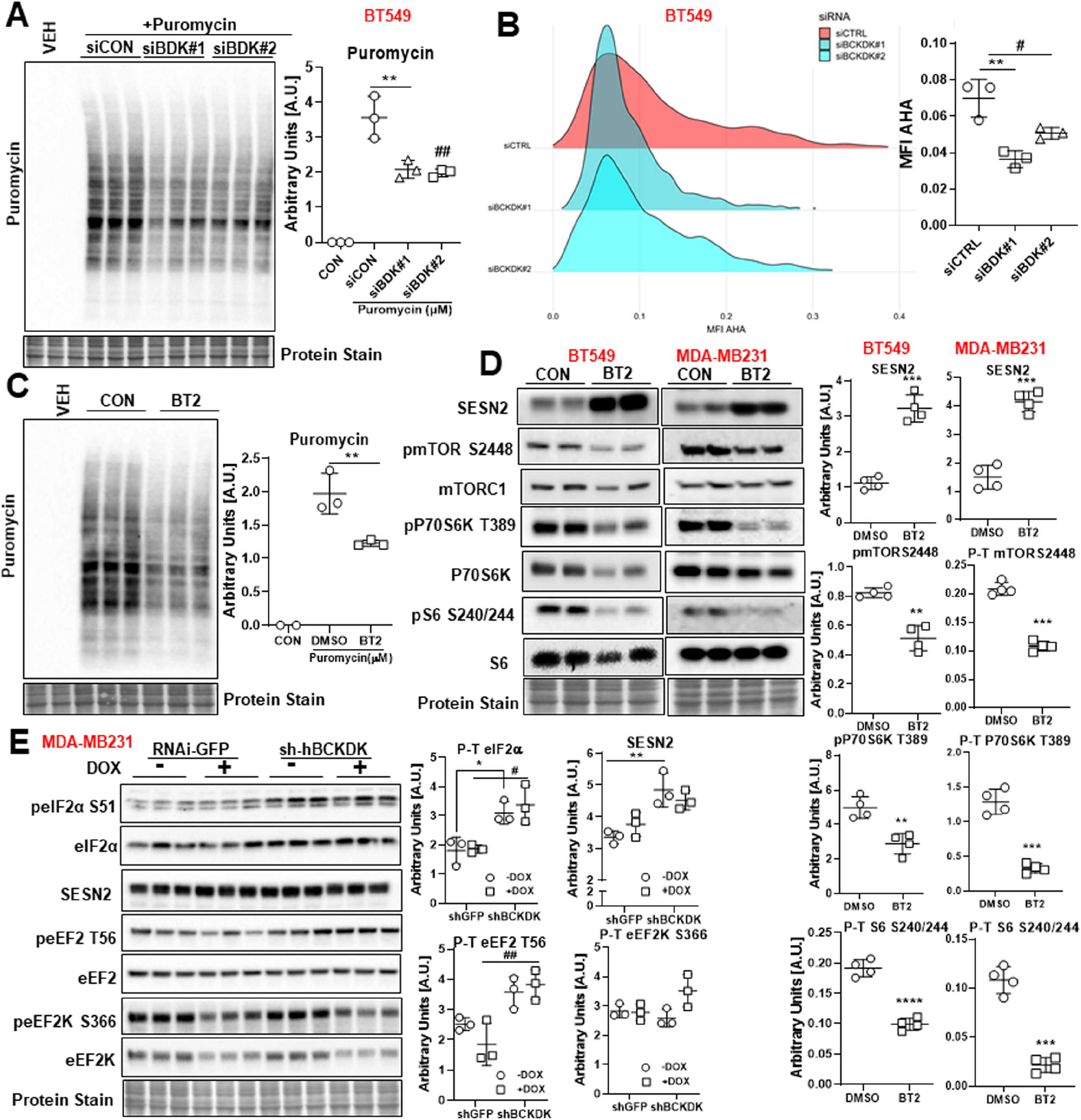
BCKDK inhibition decreases protein synthesis and mTOR signaling in TNBCs. A) BT549 transfected with siCON, siBDK#1 or siBDK#2 for 72h followed by incubation with 1μM puromycin for 30 mins. Immunoblot and densitometric analysis of puromycin incorporation. B) Fluorogram and quantification of AHA incorporation in BT549 cells transfected with siCON, siBDK#1 or siBDK#2. C) Immunoblot and densitometric analysis of puromycin incorporation in BT549 cells treated with 500μM BT2 for 20h followed by incubation with 1μM puromycin for 30 mins. The graph represents mean ± S.D., n=3, *p<0.05 was performed using Student’s t-test. D) Immunoblot and densitometric analysis of SESN2, total and phosphorylated mTOR S2448, total and phosphorylated P70S6K T389 and total to phosphorylated S6 S240/244 in BT549 and MDA-MB231 cells treated with 500μM BT2 for 20h. Data presented as mean ± S.D. Statistical analysis was performed using a was performed using Student’s t-test; *p <0.05, **p < 0.01, **** p <0.0001 as indicated. Immunoblot and densitometric analysis of total and phosphorylated eIF2α S51, Sesn2, total and phosphorylated eEF2 T56, total and phosphorylated eEF2K S266, total and phosphorylated AKT S473 and AKT T308 in MDA-MB231 cells transduced with shGFP or shBCKDK for 48h followed by 2μM DOX or DMSO treatment for 18h. Quantifications are from three independent experiments. Data presented as mean ± S.D. Statistical analysis was performed using a two-way ANOVA followed by a Tukey’s multiple comparison test; *p <0.05, **p < 0.01, **** p <0.0001 as indicated.

### Silencing BCKDK reduces protein synthesis in TNBCs

Previous studies have demonstrated that induction of ER stress and mitochondrial dysfunction^20^, induces apoptosis^21^ and inhibits protein translation and synthesis. As measured by puromycin incorporation, protein synthesis was reduced in BCKDK depleted BT549 cells (Fig 6A). Similarly, azidohomoalaine (AHA) labelled nascent protein levels were decreased upon BCKDK silencing (Fig 6B). A similar reduction in puromycin incorporation was observed in BT549 cells treated with BT2 (Fig 6C). In both BT549 and MDA-MB231 cells, BT2 treatment resulted in a robust increase in sestrin 2 (SESN2) content (Fig 6D), a negative regulator of mammalian target of rapamycin (mTOR) and protein synthesis^22^. Consistently, mTOR signaling was suppressed in BT2 treated cells as observed by reduced phosphorylation on mTOR S2448, P70S6K T389 and S6 S240/244 (Fig 6D). Similar increases in SESN2 levels were observed in MDA-MB231 cells adenovirally transduced with shBCKDK in the presence or absence of DOX (Fig 6E). Further, BCKDK inhibition in the presence or absence of DOX was accompanied by increased inactivating phosphorylation of eIF2α at S51 (Fig 6E), suggesting inhibited protein translation. BCKDK inhibition also increased phosphorylation of the elongation factor eEF2 at T56 (Fig 6E), which also correlated with suppressed protein translation.

## Discussion

TNBCs are a heterogeneous breast cancer subtype with early relapse rates, poor prognosis and limited therapeutic options^1^. Sustaining oncogenesis in TNBCs is dependent on BCAAs which contribute to protein biosynthesis, nitrogen homeostasis, epigenetic regulation, redox balance, immune response and tumor surveillance, growth and metastasis^23,24^. Numerous types of human cancers exhibit significant reprogramming of the BCAA metabolic pathway^11^. Nevertheless, the contribution of BCAA catabolic dysregulation in TNBC pathology, cell function and chemoresistance merit investigation. In the current study, we observed consistently suppressed expression of BCAA catabolic enzymes along with increased mRNA expression of the regulatory kinase, BCKDK, in two TNBC cell lines. Transcriptome analysis demonstrated that genes involved in caspase-dependent apoptosis and mitochondrial function were differentially enriched in MDA-MB231 cells with BCKDK silencing. BCKDK inhibition also sensitized TNBCs to DOX-induced apoptosis, mitochondrial ATP insufficiency and suppression of mTOR dependent protein translation by SESN2. For the first time, our data proposes a novel role of BCKDK in TNBC survival, proliferation, and DOX chemosensitivity.

Elevated intratumoral BCAAs are reported in several cancers, including breast cancer^14^, PDAC^25,26^, HCC^27^, and NSCLC^28^, while increased BCKAs secretion is observed in glioblastomas and PDACs^8,9^. However, not only BCAA metabolites but also BCAA catabolizing enzymes have recently observed to influence cancer outcomes^11^. Indeed, upregulation of BCAT1 is reported in glioblastomas, HCC, leukemias, osteosarcomas, ovarian and endometrial cancer^27,29,11^. BCAT2 is upregulated in PDAC^25,26^, luminal type breast cancer^14^, NSCLC^28^, while it is downregulated in HCC^27^. BCKDH is upregulated in leukemia and luminal type breast cancer^30,14^ and suppressed in HCC and PDAC^27,28^. Concomitantly, BCKDK expression was increased in HCC however, more distal enzymes of BCAA catabolism, ACADS and ACADSB were downregulated^27^. Our data are in agreement with these findings demonstrating dysregulated BCAA catabolic enzyme expression in MDA-MB231 and BT549 cells, highlighting the involvement of BCAA catabolizing enzymes and metabolites in tumorigenesis.

Two studies have recently identified different phosphorylation sites on BCKDK that mediates cancer proliferation, signaling and metastasis^12,13^. Notably, stabilizing BCKDK by Src promoted migration, invasion and metastasis in colorectal cancers^12^. Our data in TNBCs demonstrate that inhibiting BCKDK impairs TNBC proliferation and sensitizes TNBCs to DOX toxicity. We propose that the metabolic effects of BCKDK are likely an outcome of altered BCAA-BCKA flux towards oxidation and away from protein synthesis. Indeed, increased intracellular BCAA accumulation upon BCKDHA deletion resulted in increased cell proliferation in an immortalized hepatocyte cell line^27^. Moreover, increased extracellular secretion of BCKAs in a BCAT1 dependent manner supports cell proliferation in PDACs^8^ and survival of glioblastoma cells by evading immune surveillance^9^. Our study also demonstrated that pharmacological inhibition of BCKDK was sufficient to reduce both intracellular and secreted BCKAs in MDA-MB231 cells, plausibly contributing to reduced cell proliferation. Since TNBCs are classically treated with combination therapy involving anthracyclines^2^, such as DOX, it is plausible that targeting BCKDK might trigger a metabolic vulnerability rendering TNBCs susceptible to DOX toxicity. BCKDK is recently reported to act as a regulatory, upstream kinase of MEK^31^ and ERK1/2^13^, drivers of cell proliferation. Prior studies have inferred that BCKDK stabilization is an alternative mechanism to activate RAS/RAF/MEK/ERK signaling and confer drug resistance^31^. It remains to be determined whether the accumulation of BCAA metabolites, such as BCKAs, activates oncogenes and drives growth factor signaling in TNBCs. We theorized whether BCKDK silencing with concomitant DOX treatment potentiates BCKA oxidation reducing intracellular BCKAs compromising growth and increasing cell death. In our study, we observed that when TNBCs were exposed to DOX, BCKDK expression was significantly downregulated. Moreover, silencing BCKDK markedly exacerbated cytotoxic effects of DOX in a caspase 3/7 dependent manner and reduced cell proliferation. Although the mechanism by which BCKDK silencing triggers apoptosis induction remains unclear, we theorize that perturbations in protein synthesis and altered mitochondrial metabolism might precede or follow apoptotic events following BCKDK inhibition.

Dysregulation of protein translation and anomalous energy metabolism are characteristic of cancers, processes that are sensitive to the effects of chemotherapeutic agents^32,33^. Chemotherapeutics such as DOX can accumulate within the mitochondria resulting in mitochondrial dysfunction and energy stress by disrupting the electron transport chain (ETC) and reducing mitochondrial oxidative capacity^19^. Indeed, a decline in ATP synthesis is associated with mitochondrial accumulation of DOX^34^. We found that BCKDK inhibition markedly reduced ATP production and oxygen consumption in MDA-MB231 cells but did not exacerbate DOX mediated suppression of respiration or energy production. Notably, BCKDK deficient fibroblasts show reduced intracellular ATP and respiration, with corresponding increase in ROS production and mitochondrial hyperfusion^35^. Interestingly, targeting BCKDHA in PDACs did not affect mitochondrial metabolism or oxygen consumption^26^, suggesting the effects of BCKDK on mitochondrial function and metabolism might be independent of BCKDHA. Moreover, our transcriptomics data in BCKDK depleted cells identified reduced expression of several mitochondrial genes associated with complex formation, glycine metabolism and cholesterol trafficking. However, it remains to be determined whether targeting BCKDK in TNBCs impairs mitochondrial function due to altered mitochondrial biogenesis, mitophagy mitochondrial DOX accumulation or because of reduced entry of BCAA derived TCA cycle intermediates.

Changes in the metabolic microenvironment within TNBCs hyperactivates signaling pathways of protein translation and biosynthesis, enabling uncontrolled growth and survival. Protein synthetic rates are significantly reduced following apoptosis induction, where caspase-dependent proteolytic cleavage and altered phosphorylation states of translation initiation factors (eIF4GI, eIF4GII, eIF4E, eIF4B and eIF2α) influence cancer progression^21^,^36^. We found that BCKDK inhibition decreased protein synthesis. Consistent with this finding, upon BCKDK depletion we also observed significant increase in the inhibitory phosphorylation of the elongation factor, eEF2 at Thr 56. eEF2 phosphorylation impedes translocation of peptidyl-tRNA from the A site to the P site on the ribosome. High intracellular Ca^2+^ levels increase phosphorylation of eEF2^37^. Our transcriptomic data in BCKDK depleted cells revealed enrichment of the GO-term related to NFAT-calcineurin signaling cascade, which in turn is activated by high intracellular Ca^2+^ levels, a likely trigger for increased eEF2 phosphorylation. eEF2 is downstream of the AMPK-mTORC1 axis where it is inhibited by AMPK and activated by mTORC1. Our data in both TNBC cell types showed impaired mTORC1 signaling upon BCKDK inhibition and significant upregulation of SESN2, the negative regulator of mTORC1. The suppressed activation of mTORC1 signaling can explain the inactivation of eEF2. Additionally, reduction in protein synthesis following BCKDK inhibition was associated with increased inhibitory phosphorylation of eIF2α at Ser 51. eIF2α phosphorylation increases in response to stress and cancer^36^. eIF2α phosphorylation at Ser 51 inhibits the eIF4B catalyzed guanine nucleotide exchange function of eIF2 that prevents the formation of the 43S preinitiation complex^21^. It is unclear whether a decline in mTORC1 driven protein synthesis induces apoptosis^38^ or increased apoptosis suppress global translation or there is a diminished amino acid availability in TNBCs with inhibited BCKDK.

In conclusion, our study highlights BCKDK as a target for potentiating cytotoxic effects of DOX in TNBCs. This study demonstrates that BCKDK inhibition alone is sufficient to induce ATP insufficiency and decrease protein translation, caspase activation processes rendering TNBCs susceptible to cell death. Augmented BCKA levels due to BCKDK silencing can be fated to either get reaminated to BCAAs or get oxidised. Since BCAAs (majorly leucine) is recognised as a key amino acid to activate mTOR, the former fate for BCKAs is unlikely in our study as mTOR signaling is suppressed. However, changes in mTOR signaling may or may not be dependent on changes in BCAA levels. Further studies are required to identify whether BCKDK’s role in TNBCs are mediated via increased BCKA oxidation or is independent of the changes in intracellular BCKA flux. Moreover, it needs to be determined how BCKDK silencing triggers the activation of caspases and the apoptotic pathway and if p53 mediates these effects. DOX’s genotoxic stress also involves DNA intercalation and double-strand breaks; whether BCKDK silencing confers potentiated sensitivity to DOX by incrementing DNA damage remains to be explored. Since BCKDK was recently identified as a cytosolic kinase regulating hepatic lipid accumulation^39^, future studies will involve the identification of other metabolic pathways that co-operate with BCAA metabolism to promote tumor growth or alternatively compensate for sustaining cell proliferation when BCAA metabolism is inhibited.

## Materials and Methods

### Cell lines and culture conditions

MCF10A cells (CRL-10317, US) and BT549 cells (HTB-122) were obtained from ATCC. MDA-MB231 cells were a gift from Dr. G. Robichaud (Université de Moncton). The cells were cultured according to our previously published studies^3^. All the cell lines were maintained at 37°C in a humidified atmosphere of 5% CO_2_. The cell lines have not been genetically authenticated. For the BT2 experiments, cells were pretreated for 20h with the desired concentration of BT2 (Sigma and Matrix Scientific). BT2 from Sigma was dissolved in cremaphore solution, generously gifted by Ayappan Subbiah (Sevengenes Inc.).

### Transfections and viral transductions

Adenoviral vector expressing BCKDK shRNA (shADV-225358) and the control vector for scrambled shRNA GFP (Cat# 1122) were obtained from Vector Biolabs (USA). Adenoviral infection of cells was done 24h post-plating, and the multiplicity of infection (MOI) was kept constant between the control and experimental constructs. siRNA knockdown of BCKDK was performed using Ambion silencer select siRNA oligonucleotides (Thermo-Fisher Scientific Cat# 4390824). The siRNAs used in this paper were siBCKDK#1: #s20126, siBCKDK#2: #s20127, siRNA negative control (Cat# 4390844). Transfection was achieved using Lipofectamine RNAiMAX (Invitrogen) following the manufacturer’s instructions with a concentration of 10 nM of siRNA per plate.

### Cell lysate processing and immunoblotting

Cell pellets were sonicated in ice-cold lysis buffer (containing 20 mM Tris-HCl, pH 7.4, 5 mM EDTA, 10 mM Na_4_P_2_O_7_ (Calbiochem), 100 mM NaF, 1% Nonidet P-40, 2 mM Na_3_VO_4_, protease inhibitor (10 ml/ml; Sigma), and phosphatase inhibitor (10 ml/ml; Calbiochem) and centrifuged at 16,000g for 15 min. Protein concentration determination, immunoblotting and development of immunoblots were performed as described previously^40^. List of primary antibodies is described in Table S1. Densitometric analysis was performed using Image laboratory software (Bio-Rad), and the quantifications were normalized by total protein loading using GraphPad software (Clarivate).

### qPCR analysis

mRNA levels of BCAA-catabolizing enzymes were determined using qPCR by employing validated optimal reference gene pairs. Primer information of the target and reference genes are provided in Table S2. RNA isolation, quality control, cDNA synthesis and qPCR analysis were performed as described previously^41^.

### Trypan blue exclusion and metabolic activity assays

For cell viability, 1 × 10^4^ cells were plated in 24-well plates in triplicates. Total number of viable cells were calculated using the trypan blue exclusion assay. For measuring metabolic activity, cells were plated in 96-well plates and treated with DOX for 18h followed by incubated in drug-free media for 48h. Metabolic activity of viable cells were measured by incubating the cells with Presto Blue (Thermo-Fischer Scientific) for 3h at 37°C. Fluorescent intensity was read at Ex/Em= 560/590 on a microplate fluorometer (Synergy H4).

### CCK8 assay

Cell Counting Kit-8 (CCK-8; Sigma Aldrich) was used to measure cell viability according to the manufacturer’s protocol. Briefly, 1 x 10^5^ MDA-MB231 cells were seeded in 96-well plates transfected with the siRNAs for different time points followed by DOX treatment for 18h. Following the treatments, the cells were incubated in CCK-8 solution at 37 ℃ for 4h. Synergy H4 microplate reader was used to measure the absorbance at 450nm.

### Extracellular flux analyzer studies

MDA-MB231 or MCF10A cells were plated at a density of 30,000 cells/per well. Mitostress assay was performed according to Khan et al.^42^ using the Seahorse XFe24 analyzer (Agilent Technologies Inc. CA, USA). Briefly, cells were incubated with XF Assay medium (with 20 mM glucose, 1 mM sodium pyruvate and 1 mM glutamate, without sodium bicarbonate) for 1 h. 1 μM of oligomycin, 1 μM of FCCP [carbonyl cyanide‐4‐ (trifluoromethoxy)phenylhydrazone], and 1 μM each of rote‐ none and antimycin A was injected over 100 min to measure the different stages of respiration. The assay was normalized with protein and analyzed with the XFe 2.0.0 software (Agilent Technologies Inc. CA, USA).

### Apoptosis array

Expression of 43 apoptosis-associated proteins in duplicates were determined by the Human apoptosis antibody array (ab134001, Abcam). Membranes were incubated with 250μg of cell extracts, immunoblotted and developed using the chemidoc (BioRad) according to manufacturers’ instructions. Densitometric analysis was performed using Image laboratory software (Bio-Rad).

### Lactate dehydrogenase cytotoxicity assay

Lactate dehydrogenase (LDH) cytotoxicity assay was performed using the assay kit (Invitrogen, C20301) according to the manufacturer’s instructions. Briefly, 1×10^4^ cells were plated in a 96 well plate for experimental conditions as well as spontaneous and maximum LDH activity controls. On the day of the assay, 10 μl of 10X lysis buffer was added to wells serving as the maximum LDH activity controls and incubated at 37°C for 45 min. The reaction mixture was added to the collected media in a 1:1 ration and incubated at RT for 30min in the dark, followed by adding a stop solution. The absorbance was measured at 490 nm, and 680 nm and the assay was normalized by protein concentration.

### Caspase 3 activity assay

Caspase activity assay (Caspase-3 Activity Assay Kit #5723, Cell Signaling Technology, Danvers, MA, USA) was performed according the manufacturer’s instructions. Briefly, 1×10^4^ cells were plated in a 96 well plate and were harvested using ice-cold lysis buffer. Lysates from two wells within the same treatment group were combined for all treatment groups and transferred to a black plate. Fluorescence was measured immediately at Ex/Em 380/440 for the 0h reading. A second reading was taken after 1h incubation in the dark at 37°C. The assay was normalized by protein concentration.

### AHA incorporation assay

AHA incorporation analysis was performed using the Click-iT AHA Alexa Fluor 488 Imaging Kit (Thermo-Fisher Scientific) according to the manufacturer’s instructions. Briefly, cells on coverslips were pulse labelled with AHA for 30 min in methionine/cysteine free DMEM media without serum before performing the click labeling. Coverslips were mounted in Prolong anti-fade mounting media (Thermo-Fisher Scientific) and imaged with a Zeiss Axio Observer Z1 equipped with an Apotome.2 structural illumination unit using a 20x Plan-Apochromat objective (NA: 0.8, air). Images were processed for analysis in ImageJ, and image processing was identical for each image set. Images were analyzed in Cell Profiler, and data are represented as the mean ± S.D. per cell of three experiments.

### RNA-seq analysis and bioinformatics

MDA-MB231 cells were transfected with either of two siRNA’s targeting BCKDK or a scrambled siRNA and cultured for 48 h before cells were harvested and RNA extracted using the Qiagen RNeasy mini kit according to the manufacturer’s instructions. RNA-Seq was conducted by McGill University and the Genome Quebec Innovation Center (Montreal, Canada) with the Illumina NovaSeq 6000 S2 PE100 — 50 M platform. Analysis of the transcriptomic data was performed according to our previous published study^3^. Testing for differential expression was conducted only on genes with an average estimated count of greater than 0.3. Genes were considered significantly differentially expressed if the adjusted P-value was less than 0.01 for both siBCKDK#1 and siBCKDK#2 groups, and the fold change was occurring in the same direction. RNA-Seq data is desposited in NCBI’s Gene Expression Omnibus under the assecession number GSE163297.

### SunSET method

Protein synthesis was measured in vitro by the SunSET assay as described previously ^40^. Briefly, BT549 cells were either transfected with siBCKDK #1 or #2 and siCon for 48h or with 500μM BT2 for 20h followed by treatment with 1 μM puromycin dihydrochloride (P8833, Sigma) for 30 min followed by a 30min chase with complete media. Puromycin incorporation was detected by Western blotting using the monoclonal puromycin antibody.

### BCKA measurements

For BCKA measurements, cells were incubated with serum-free DMEM low glucose, leucine-free medium (D9443, Sigma) for 24h or during the duration of the treatment.

#### Secreted BCKA extraction

20 μl of the media and 120 μl of internal standard (4 mg/ml in H2O) containing leucine-d3 (CDN Isotopes), 40 μl of MilliQ water, 60 μl of 4 M perchloric acid (VWR) were combined and vortexed. Proteins were precipitated in two sequential steps, followed by centrifugation at 13,000 rpm for 15 min at 4 °C. Supernatants collected from both steps were combined for measuring BCKAs. Intracellular BCKA extraction, BCKA derivatization and quantification were done according to our previously published study^40^.

### Statistical analysis

Data are expressed as mean±S.D. unless otherwise indicated. Statistical analyses were conducted using Prism software (GraphPad, La Jolla, CA, USA). Comparisons between multiple groups were performed using one-way or two-way analysis of variance followed by a Tukey post hoc test, as appropriate. Data sets with two groups were analyzed using a two-tailed student’s t-test. p values of 0.05 were considered statistically significant.

## Abbreviations

Acadsb: Acyl-CoA dehydrogenase short branched chain
ATM: ataxia-telangiectasia mutated
BCAA: branched-chain amino acid
BCKA: branched-chain keto acid
BCAT1: branched-chain aminotransferase
BCKDHA: Branched chain keto acid dehydrogenase E1 alpha polypeptide
BCKDHB: Branched chain keto acid dehydrogenase E1 subunit beta
BCKDK: branched-chain keto acid dehydrogenase kinase
BT2: 3,6-dichlorobenzo[b]thiophene-2-carboxylic acid
eEF2: eukaryotic elongation factor 2
eEF2K: eukaryotic elongation factor 2 kinase
eIF2α: eukaryotic translation initiation factor 2A
HADHA: Hydroxyacyl-CoA dehydrogenase trifunctional multienzyme complex subunit alpha
HIBCH: 3-hydroxyisobutyryl-CoA hydrolase
HIBADH: 3-hydroxyisobutyrate dehydrogenase
HSP90AB1: Heat shock protein HSP 90-beta
KLF15: Kruppel like factor 15
mTORC1: mammalian target of rapamycin
PARP: poly (ADP-ribose) polymerase
PPM1K: Protein phosphatase Mg2+/Mn2+ 1K
SESN2: sestrin2.

## Declaration

### Funding & Acknowledgement

This work was supported by Natural Sciences and Engineering Research Council of Canada (RGPIN-2020-05906), Diabetes Canada (NOD_OG-3-15-5037-TP & NOD_SC-5-16-5054-TP) and New Brunswick Health Research Foundation grants to T.P.; D.B., was funded by postdoctoral fellowships from New Brunswick Health Research Foundation and Dalhousie Medicine New Brunswick.

### Data availability

All data and materials used in the current study are available from the corresponding author upon request. The datasets geberated and/or analysed during the current study are available in NCBI’s NCBI’s Gene Expression Omnibus.

Accession ID: GSE163297

Databank URL: https://www.ncbi.nlm.nih.gov/geo/query/acc.cgi?acc=GSE163297

## SUPPORTING INFORMATION

**Figure S1.**
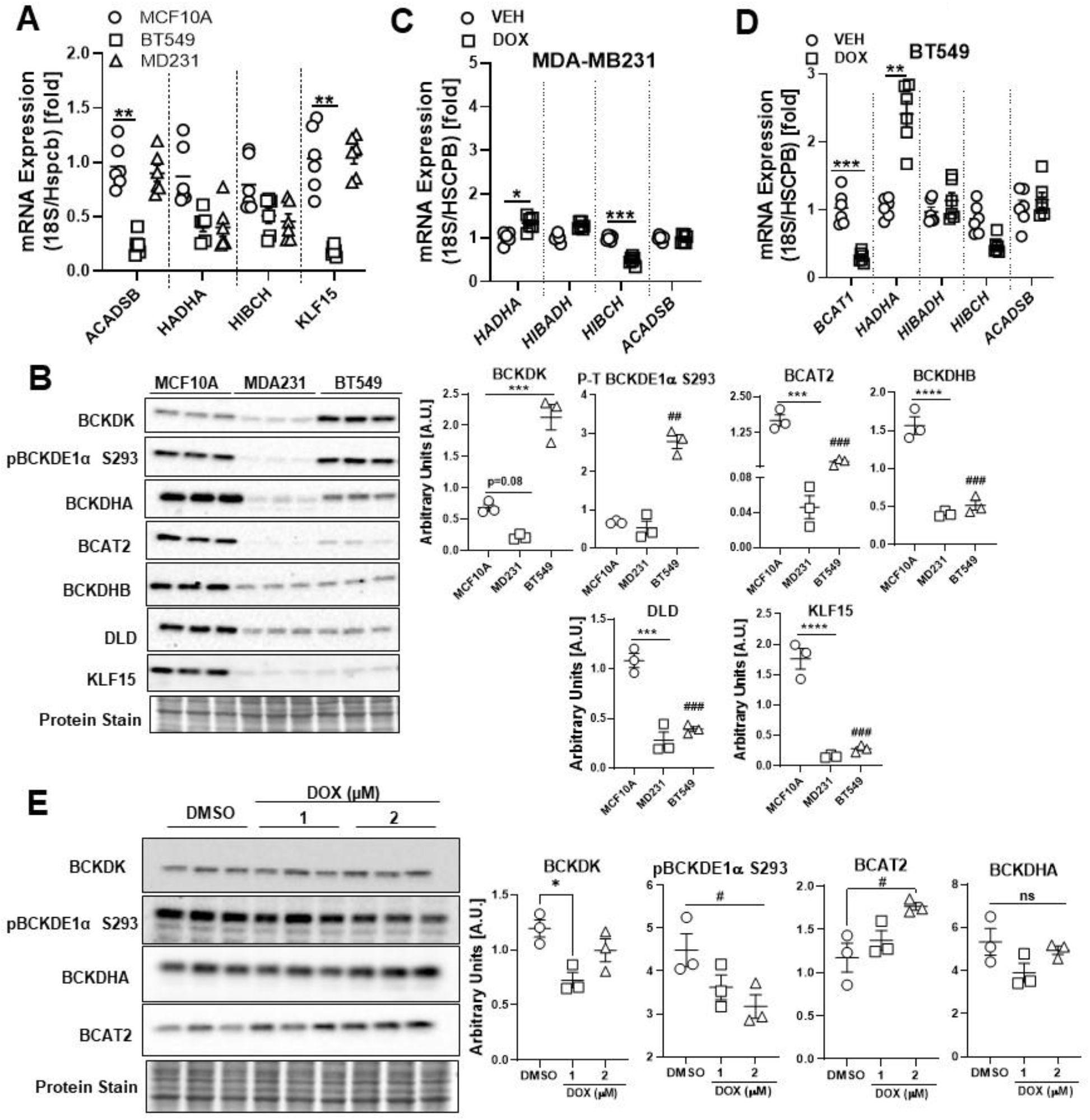
BCAA degradation enzyme expression are suppressed in TNBCs and can be revived by DOX treatment. A) Quantification of *ACADSB*, *HADHA*, *HIBCH*, and *KLF15* BCAA catabolic enzyme mRNA expression corrected to 18S/HSPCB reference genes in MCF10A, BT549 and MDA-MB231 cells. B) Immunoblot and densitometric analysis of BCKDK, total and phosphorylated BCKDHA E1α Ser 293, BCAT2, BCKDHB, DLD and KLF15 in MCF10A, MDA-MB231 and BT549 cells. C-D) BCAA catabolic enzyme expression in TNBCs treated with 2μM DOX for 18h. Quantification of *BCAT1*, *ACADSB*, *HADHA*, *HIBCH*, mRNA expression corrected to 18S/HSPCB reference genes in MDA-MB231 (C) and BT549 (D) cells. E) Immunoblot and densitometric analysis of BCKDK, total and phosphorylated BCKDHA E1α Ser 293 and BCAT2 in BT549 cells treated with 1μM and 2μM DOX for 18h. Data presented as mean ± S.D. Statistical analysis was performed using a two-way ANOVA followed by a Tukey’s multiple comparison test; *p <0.05, **p < 0.01, **** p <0.0001 as indicated.

**Figure S2.**
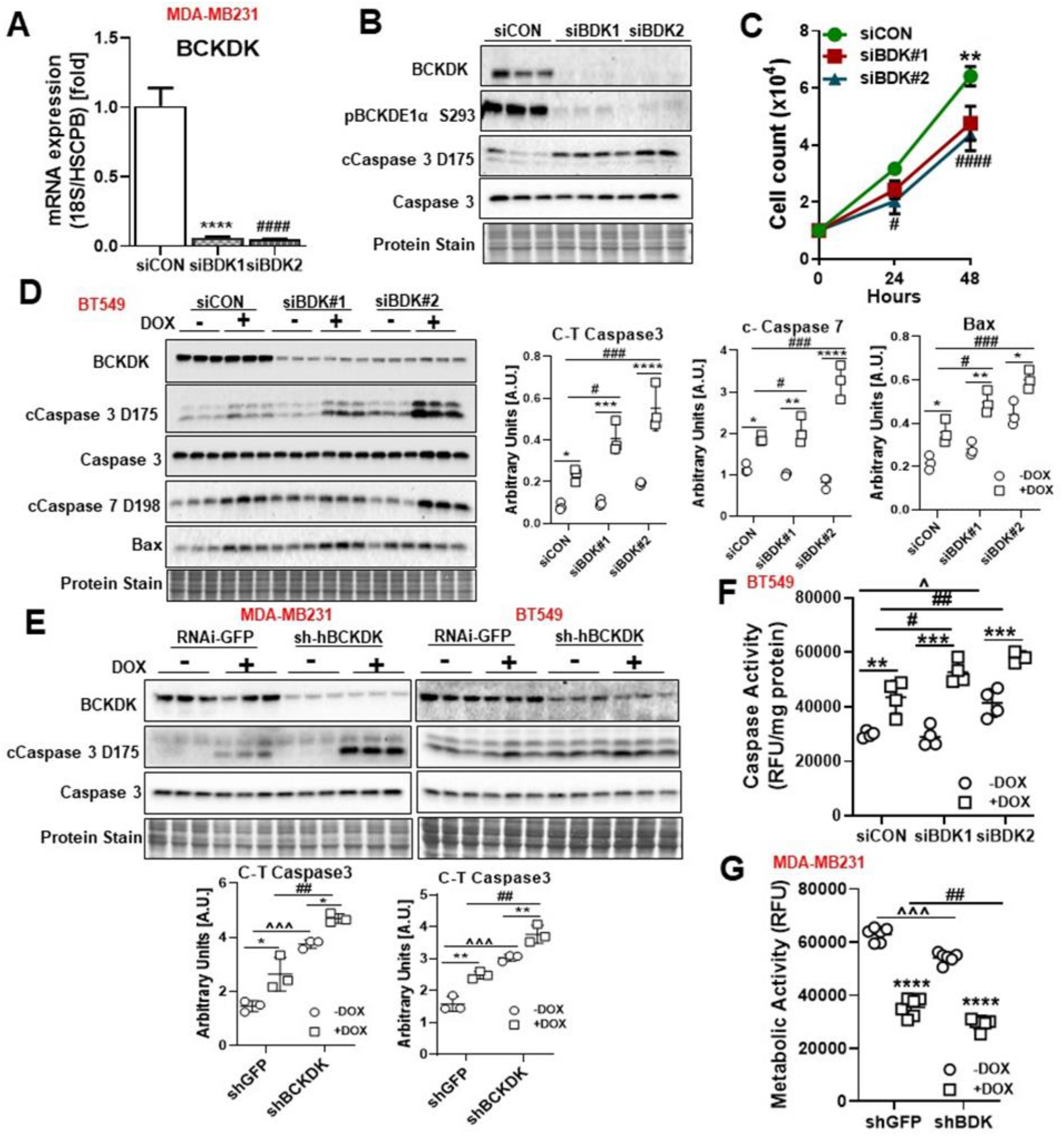
BCKDK knockdown increases cell death and reduces proliferation and potentiates DOX’s effects in TNBCs. mRNA quantification of *BCKDK* (A) and protein expression (B) of BCKDK and cleaved and total caspase 3 expression in MDA-MB231 cells transfected with siRNA targeting BCKDK at exon 4 (siBDK#1) and exon 5 (siBDK#2). C) Cell counts at 0, 24 and 48h of MDA-MB231 cells transfected with siCON, siBDK#1 or siBDK#2. D) Immunoblot and densitometric analysis of BCKDK, total and cleaved Caspase 3, cleaved Caspase 7, total and Bax in BT549 cells transfected with siCON, siBDK#1 or siBDK#2 for 72h followed by 2μM DOX or DMSO for 18h. E) Immunoblot and densitometric analysis of total and cleaved Caspase 3 in MDA-MB231 and BT549 cells transduced with either shGFP or shBCKDK for 48h followed by 2μM DOX or DMSO for 18h. F) BT549 cells were transfected with siCON, siBDK#1 or siBDK#2 for 72h followed by 2μM DOX or DMSO treatment for 18h and analyzed for Caspase 3 activity. G) Metabolic viability of MDA-MB231 cells transduced with either shGFP or shBCKDK and treated with 2μM DOX or DMSO for 18 h plus 48 h in drug-free medium. Data presented as mean ± S.D. Statistical analysis was performed using a two-way ANOVA followed by a Tukey’s multiple comparison test; *p <0.05, **p < 0.01, **** p <0.0001 as indicated.

**Figure S3.**
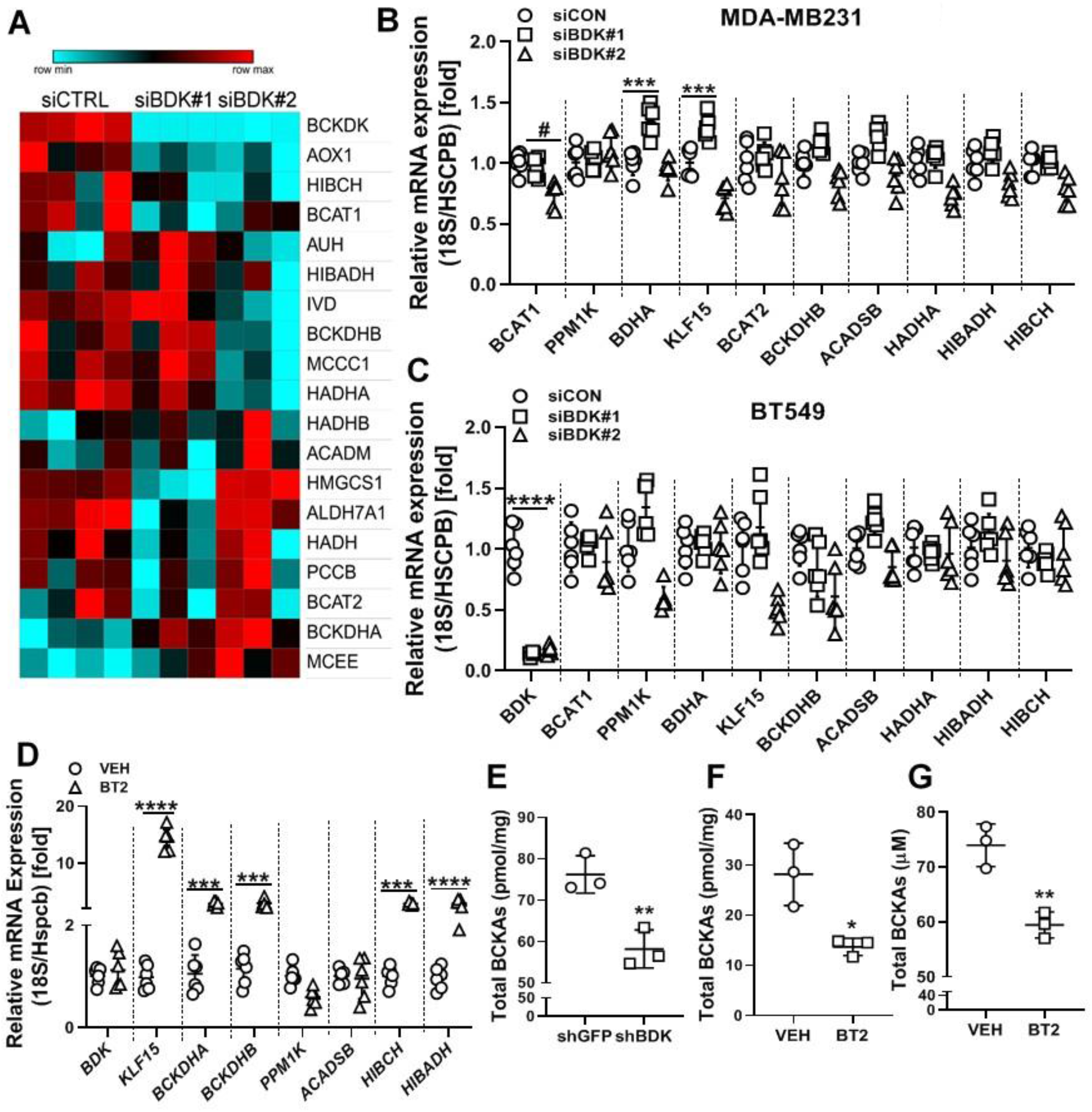
BT2 augments mRNA expression of BCAA catabolic genes and reduces BCKA accumulation and secretion. A) Heatmap for the genes involved in BCAA catabolism which were differentially regulated by BCKDK knockdown. B-C) Quantification of BCKDK, BCKDHA, PPM1K, BCKDHB, BCAT2, BCAT1, ACADSB, HADHA, HIBCH, HIBADH and KLF15 mRNA expression corrected to 18S/HSPCB reference genes in B) MDA-MB231 and C) BT549 cells transfected with siCON, siBDK#1 or siBDK#2 for 72h. D) Quantification of BCKDK, BCKDHA, PPM1K, BCKDHB, ACADSB, HIBCH, HIBADH and KLF15 mRNA expression corrected to 18S/HSPCB reference genes in BT549 cells treated with 500μM BT2 for 20h. Statistical analysis was performed using a two-way ANOVA followed by a Tukey’s multiple comparison test; *p <0.05, **p < 0.01, **** p <0.0001 as indicated. E) UPLC MS/MS analysis of intracellular BCKAs in MDA-MB231 cells transduced with shBCKDK and shGFP for 48h. Measurement of intracellular (F) and secreted BCKAs in the media (G) by UPLC MS/MS in MDA-MB231 cells treated with 500μM BT2 for 20h. Data presented as mean ± S.D. Statistical analysis was performed using a Student’s t-test; *p <0.05, **p < 0.01, **** p <0.0001 as indicated.

**Figure S4.**
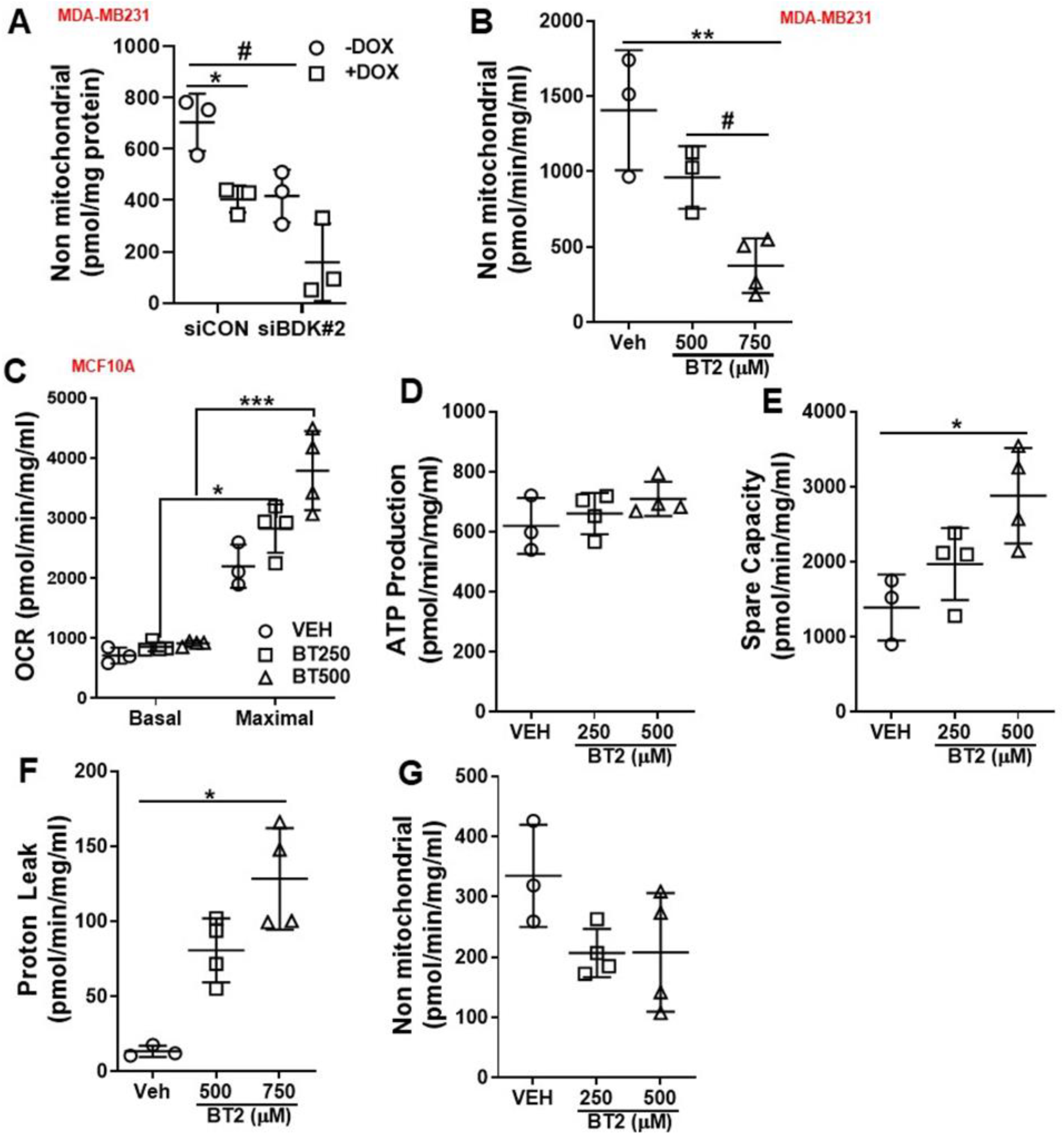
BT2 does not affect mitochondrial respiration in MCF10A cells. A-B) Non mitochondrial respiration measured in MDA-MB231 cells A) transfected with siCON or siBDK#2 for 48h or B) treated with 250μM or 500μM BT2 for 20h, in the presence of 25mM glucose. C-G) Basal and maximal OCR (C), ATP production (D), spare capacity (E), proton leak (F) and non mitochondrial respiration (G) measured in MCF10A cells treated with 250μM or 500μM BT2 for 20h in the presence of 25mM glucose. Data presented as mean ± S.D. Statistical analysis was performed using a was performed using two-way ANOVA followed by a Tukey’s multiple comparison test for (A) or Student’s t-test for (B); *p <0.05, **p < 0.01, **** p <0.0001 as indicated.

**Table S1.** List of antibodies. (Fine chemicals, if noted otherwise under the Materials and Methods segment, are from Sigma.)

| **Antibodies** | **Source** | **Identifier** |
| --- | --- | --- |
| BCKDK | My Biosource | MBS 275719 |
| p-BCKDE1α S293 | Bethyl Lab | A303-567A |
| BCKDHA | My Biosource | MBS 275832 |
| BCKDHB | Santa Cruz Biotechnology Inc. | H-6; sc-374630 |
| BCAT2 | Invitrogen | PA5-21549 |
| DLD | Santa Cruz Biotechnology Inc. | G-2; sc-365977 |
| KLF15 | Novus Biologicals | NBP2-24635 |
| cCaspase 3 D175 | Cell Signaling Technologies | 9664 |
| cCaspase 7 D198 | Cell Signaling Technologies | 8438 |
| Caspase 3 | Cell Signaling Technologies | 9662 |
| cPARP D124 | Abcam | ab110315 |
| PARP | Cell Signaling Technologies | 9532 |
| p-ATM S1981 | Cell Signaling Technologies | 5883 |
| ATM | Cell Signaling Technologies | 2873 |
| Sestrin 2 | Cell Signaling Technologies | 8487S |
| p-mTOR S2448 | Cell Signaling Technologies | 5536 |
| mTORC1 | Cell Signaling Technologies | 2972 |
| p-P70S6K T389 | Cell Signaling Technologies | 9234 |
| P70S6K | Cell Signaling Technologies | 2708 |
| p-S6 S240/244 | Cell Signaling Technologies | 5364 |
| S6 | Cell Signaling Technologies | 2217 |
| p-eEF2 T56 | Cell Signaling Technologies | 2331 |
| eEF2 | Cell Signaling Technologies | 2332 |
| p-eEF2K S366 | Cell Signaling Technologies | 3691 |
| eEF2K | Cell Signaling Technologies | 3692 |
| p-eIF2α S51 | Cell Signaling Technologies | 9721 |
| eIF2α | Cell Signaling Technologies | 9722 |
| Puromycin clone 12D10 | Millipore-Sigma | MABE343 |
| Total OXPHOS cocktail | Abcam | ab110413 |
| mouse anti-rabbit IgG-HRP | Santa Cruz Biotechnology Inc. | sc-2357 |
| m-IgG kappa BP-HRP | Santa Cruz Biotechnology Inc. | sc-516102 |

**Table S2.**
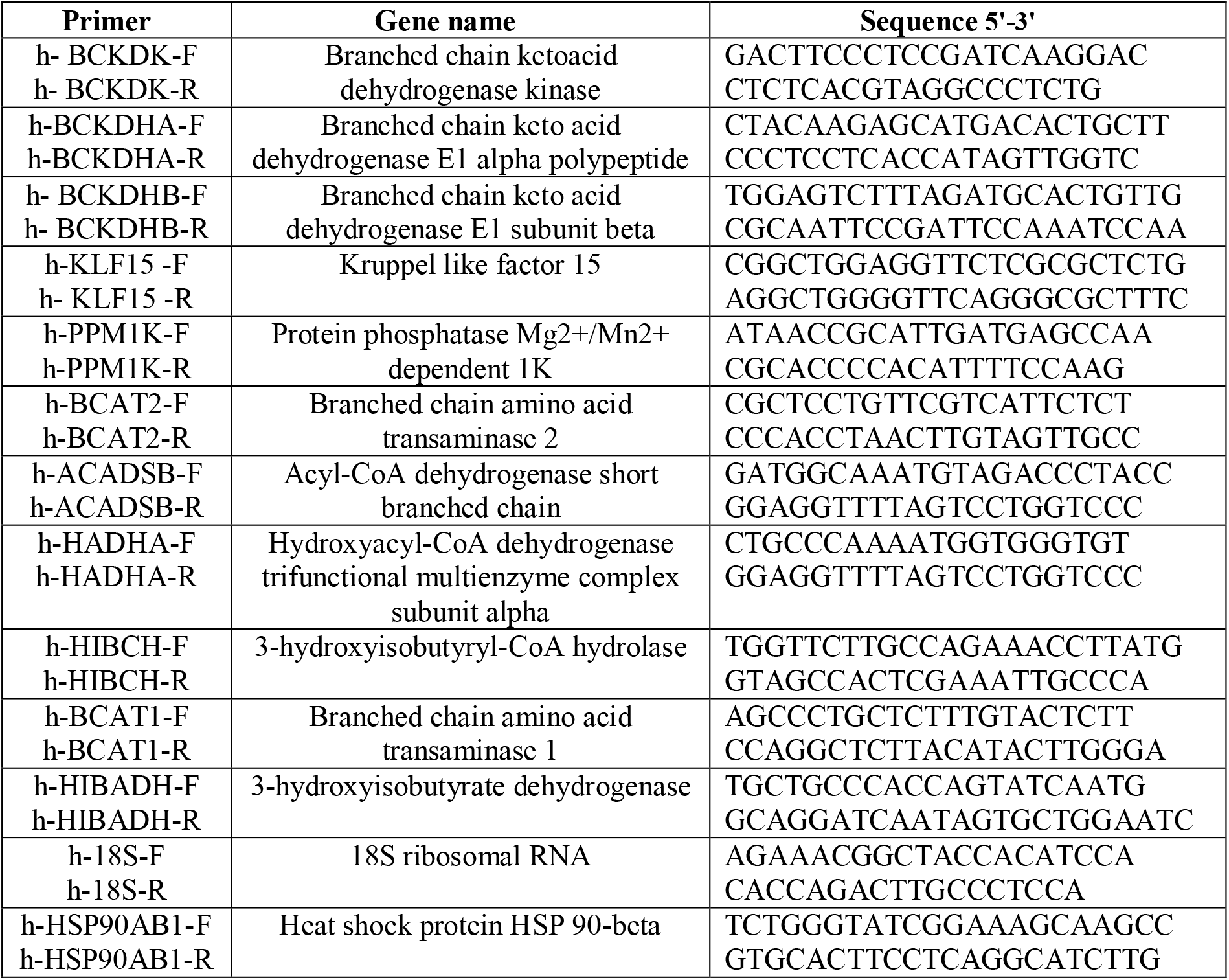
List of primers.

